# H3K27 and H3K9 Methylation Mask Potential CTCF Binding Sites to Maintain 3D Genome Integrity

**DOI:** 10.1101/2025.04.02.646724

**Authors:** Kei Fukuda, Chikako Shimura, Yoichi Shinkai

## Abstract

The three-dimensional (3D) genome structure is essential for gene regulation and various genomic functions. CTCF plays a key role in organizing Topologically Associated Domains (TADs) and promoter-enhancer loops, contributing to proper cell differentiation and development. Although CTCF binds the genome with high sequence specificity, its binding sites are dynamically regulated during development, and aberrant CTCF binding is linked to diseases such as cancer and neurological disorders, and aging. However, the mechanisms controlling CTCF binding remain unclear. Here, we investigated the role of repressive chromatin modifications in CTCF binding using H3K9 methyltransferase-deficient immortalized mouse embryonic fibroblasts (iMEFs) and H3K27 methyltransferase EZH1/2 inhibitor. We found that H3K9 and H3K27 methylation regulate CTCF binding at distinct genomic regions, and their simultaneous loss induces drastic changes in CTCF binding. These changes were associated with alterations in 3D genome architecture and gene expression, suggesting that repressive chromatin modifications preserve proper chromatin organization by preventing aberrant CTCF binding. Additionally, while CTCF binding sites repressed by H3K9 methylation were bound by CTCF in early mouse embryos, those repressed by both H3K9 and H3K27 methylation remained inaccessible, with early embryonic-specific H3K27 methylation forming at these sites. These findings implicate that H3K27 methylation prevents abnormal CTCF binding in early embryos, ensuring proper genome organization during development.

## INTRODUCTION

CCCTC-binding factor (CTCF) is a conserved transcriptional regulator composed of 11 central zinc-finger domains (ZFs). It plays a crucial role in 3D genome organization and transcriptional regulation by mediating chromatin loop formation with cohesion which acting as an insulator and contributing to the establishment of topologically associating domains (TADs) (Dixon et al. 2012; Nora et al. 2017). However, CTCF binding sites vary significantly across developmental stages and cell types, and their regulation is influenced by transcription factors and epigenomic modifications, such as DNA methylation (Monteagudo-Sanchez et al. 2024). Recent studies have revealed that SETDB1 and EHMT1/2, which are histone H3 lysine 9 (H3K9) methyltransferases, prevent ectopic CTCF binding, particularly at transposable elements (Jiang et al. 2020; Gualdrini et al. 2022; Sun et al. 2024; Tam et al. 2024). Transposable elements constitute nearly 50% of mouse genome, and some, such as SINE elements, contain CTCF-binding motifs that can occasionally function as insulators (Cournac et al. 2016; Ichiyanagi et al. 2021; Wang et al. 2024). H3K9 methylation is a key suppressor of transposable elements and may also prevent ectopic CTCF binding by blocking CTCF recognition at transposons genome-wide, thereby maintaining proper 3D genome organization. At least five H3K9 methyltransferases exist in mammals (*Setdb1, Suv3Sh1, Suv3Sh2, Ehmt1, Ehmt2*) (Allshire and Madhani 2018), and they regulate H3K9 methylation redundantly or independently, depending on genomic regions (Fukuda et al. 2021). However, due to the redundancy of H3K9 methyltransferases, their roles in CTCF binding regulation have not been fully elucidated. Furthermore, the loss of H3K9 methylation leads to epigenomic reorganization, including the redistribution of H3K27me3 mediated by the Polycomb complex (Peters et al. 2003; Walter et al. 2016; Fukuda et al. 2023). For example, H3K27me3 can spread into regions previously marked by H3K9 methylation, forming new Polycomb domains and contributing to heterochromatin maintenance following the loss of H3K9 methylation (Fukuda et al. 2023). Such epigenomic reorganization is observed not only in developmental processes such as germ cell development and early embryogenesis but also in aging and cancer (Yang et al. 2023). The Polycomb complex is known to regulate chromatin architecture through the formation of Polycomb-associated domains (PADs) and chromatin compartments (Du et al. 2020; Fukuda et al. 2023). However, its role in CTCF binding regulation and the functional implications of epigenomic reorganization in CTCF regulation remain largely unknown.

In this study, we utilize a series of H3K9 methyltransferase knockout cells, as well as a complete H3K9 methyltransferase-deficient cell line that we have successfully established (Fukuda et al. 2023). Additionally, we employ H3K27 methyltransferase EZH1/2 inhibitor to create a system in which both H3K9 and H3K27 methylation are simultaneously depleted. Using this system, we will perform CTCF ChIP-seq, RNA-seq, and Hi-C analyses to elucidate the redundant and independent roles of these repressive modifications in CTCF binding, transcription and 3D genome regulation.

## RESULTS

### H3KG and H3K27 methylation deficiency reshapes CTCF binding profiles

To investigate the function of H3K9 and H3K27 methyltransferases on CTCF binding profiles in mammalian cells, we performed CTCF Chromatin immunoprecipitation sequencing (ChIP-seq) in wild type (WT) immortalized mouse embryonic fibroblasts (iMEFs), *Setdb1* iMEFs (Kato et al. 2018), *Setdb1/Suv3Sh1/Suv3Sh2 triple* knockout (TKO) iMEFs (Fukuda et al. 2021), *Setdb1/Suv3Sh1/Suv3Sh2/Ehmt1/Ehmt2 quintuple* knockout (5KO) iMEFs (Fukuda et al. 2023), and WT, *Setdb1* KO, TKO and 5KO iMEFs treated with the EZH1/2 dual inhibitor DS3201 (hereafter referred to as DS) (Yamagishi et al. 2019). We previously demonstrated that a 7-day treatment with 1 μM DS leads to an almost complete loss of H3K27me3 in iMEFs (Fukuda et al. 2023); therefore, we applied this condition for CTCF ChIP-seq analysis in this study. CTCF ChIP-seq was performed with two biological replicates for each sample, and two clones (#14 and #55) of the 5KO iMEFs were used. Cluster analysis of the CTCF ChIP-seq data showed a high correlation between the replicates (Fig.1A). To investigate the features of CTCF binding regulated by H3K9 and H3K27 methyltransferases, we identified differentially enriched peaks (DE peaks) in each sample compared to WT using diffBind (FDR < 0.05) (Ross-Innes et al. 2012). The number of CTCF peaks obtained in each sample showed an increased CTCF peaks in DS-treated samples, especially in 5KO (Fig. 1B). In consistent with this result, the most DE peaks were observed in 5KO+DS (Fig. 1C). In samples lacking H3K9 methyltransferases (*Setdb1* KO, TKO, 5KO), the number of newly emerged CTCF peaks (increased DE peaks) in each KO cells was greater than those lost peaks (decreased DE peaks) (Fig. 1C). This suggests that H3K9 methyltransferases primarily function to prevent CTCF binding, consistent with previous analyses in *Setdb1*-deficient cells (Gualdrini et al. 2022; Sun et al. 2024; Tam et al. 2024) and *Ehmt2*-deficient cells (Jiang et al. 2020) (Fig. 1C). On the other hand, inhibition of H3K27 methyltransferases resulted in a greater number of CTCF decreased DE peaks (Fig. 1C). We noticed that treatment of each H3K9 methyltransferase-deficient cell line with DS resulted in more extensive changes in CTCF binding than the sum of CTCF changes observed in the deficient cells and those induced by DS treatment in WT cells, and this trend is particularly pronounced in 5KO cells (Fig. 1B). These results suggest that H3K27me3 has a positive effect on CTCF binding, although it is not known whether directly or indirectly, and that it also regulates CTCF binding in concert with H3K9 methylation (H3K9 methyltransferases).

**Figure 1.**
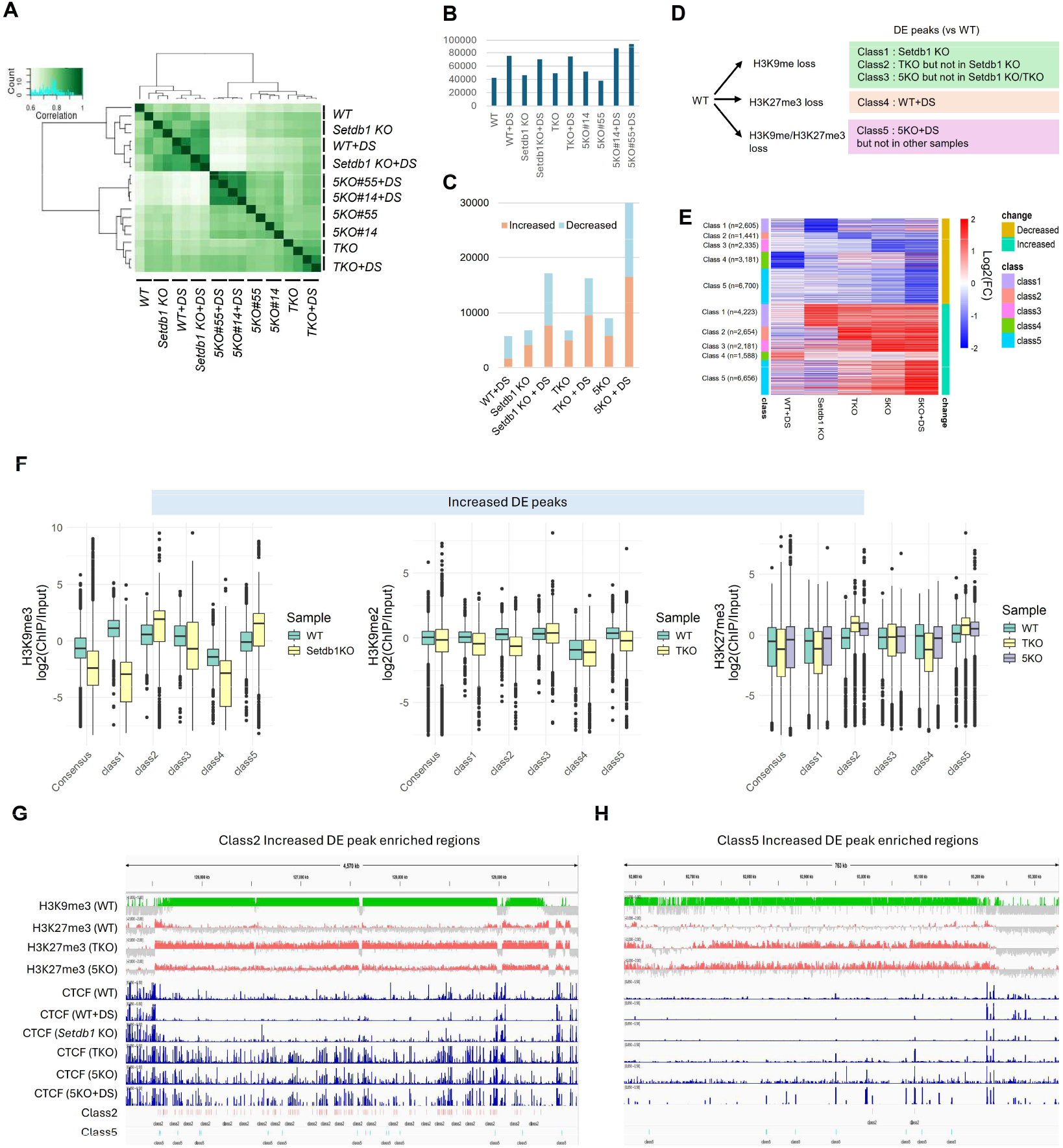
Dysregulation of CTCF binding profiles by H3K9/K27 methylation deficiency. (A) Clustering analysis of CTCF ChIP-seq data. Each sample analyzed in two biological replicates, showing a high correlation between replicates. (B) Average number of CTCF peaks in each sample. The number of peaks is averaged across replicates and displayed as a bar graph. (C) The number of differentially enriched (DE) peaks. A bar graph represents the number of CTCF accumulation increased or decreased in each sample. (D) The classification method for DE peaks. (E) Heatmap of CTCF enrichment among DE classes. (F) Epigenomic profiles at increased DE peaks. H3K9me3 enrichment in WT and Setdb1 KO iMEFs (Left), H3K9me2 enrichment in WT and TKO iMEFs (Middle), and H3K27me3 enrichment in WT, TKO, and 5KO iMEFs (Right). (G) Representative class 2 increased DE peaks enriched regions. (H) Representative class 5 increased DE peaks enriched regions.

We classified peaks with newly increased CTCF binding (Increased DE peaks) and lost peaks (Decreased DE peaks) in KO or DS-treated WT or KO iMEFs into five categories, respectively: Class 1: DE peaks observed in *Setdb1* KO (*Setdb1-dependent change*). Class 2: DE peaks observed in TKO but not in *Setdb1* KO (*Suv3Sh1/2-dependent change*). Class 3: DE peaks observed in 5KO but not in *Setdb1* KO or TKO (*Ehmt1/2-dependent change*). Class 4: DE peaks observed in WT + DS (*Ezh1/2-dependent change*). Class 5: DE peaks observed only in 5KO + DS (*Regulated by both H3K9 and K27 methyltransferases*) (Fig. 1D, E). All peaks used in *diffBind* were defined as consensus peaks, including those identified in samples other than WT iMEFs. Motif enrichment analysis of each class revealed an accumulation of CTCF-binding motifs across all classes (Supplementary Fig. S1A). Additionally, motifs such as THRB, Nr2f2, BMAL1, and NPAS were particularly enriched in “Increased” peaks observed in H3K9 methyltransferase-deficient cells, suggesting that H3K9 methyltransferases inhibit CTCF binding to motifs co-localized with these transcription factors (Supplementary Fig. S1A). Genomic distribution is largely different among classes. “Increased” Class 2, Class 3, and Class 5, are more enriched in intergenic regions (P<2.2×10^−16^) (Supplementary Fig. 1B), and the B compartments (Supplementary Fig. 1C). This is consistent with that EHMT1/2 (G9a/GLP) and SUV39H1/2 targets gene-poor intergenic regions and the B compartments (Fukuda et al. 2021), and H3K27me3 spreads into gene-poor B compartments after the loss of H3K9 methylation (Fukuda et al. 2023). “Increased” Class 2, Class 3, and Class 5 exhibit a relatively high frequency of B-to-A conversion in TKO, 5KO, and 5KO+DS, respectively, suggesting a potential association between increased CTCF binding and compartment conversion (Supplementary Fig. 1C). Unlike “Increased” Class 2, Class 3, and Class 5, “Increased” Class 1 is primarily found in the A compartment, consistent with the role of *Setdb1* in regulating H3K9 methylation mainly in the A compartment (Supplementary Fig. 1C) (Fukuda et al. 2021). Contrary to “Increased” classes, “Decreased” classes are predominantly found in genic regions, except for Class 1 (Supplementary Fig. 1B). Additionally, the compartment distribution of “Decreased” classes is the opposite of “Increased” classes, with Class 1 being enriched in the B compartment, while Class 2, 3, and 5 are more frequently found in the A compartment (Supplementary Fig. 1C).

Next, we investigated the relationship between changes in CTCF binding and chromatin modifications. “Increased” Class 1, which is regulated by *Setdb1*, exhibited a significant decrease in H3K9me3 in *Setdb1* KO (Fig. 1F). In contrast, “Increased” Class 2, which is regulated by *Suv3Sh1/2*, did not. This indicates that both H3K9me3 and CTCF binding in “Increased” Class 1 are regulated by *Setdb1*, whereas in “Increased” Class 2, both are controlled by *Suv3Sh1/2* (Fig. 1F). Regarding H3K9me2, it decreased in TKO in “Increased” Class 1 and Class 2, whereas it was maintained in “Increased” Class 3 (Fig. 1F). This suggests that both H3K9me2 and CTCF binding in “Increased” Class 3 are regulated by *Ehmt1/2*. Therefore, the differences among “Increased” Class 1, Class 2, and Class 3 can be explained by the distinct genomic regions where each H3K9 methyltransferase regulates H3K9 methylation. For H3K27me3, “Increased” Class 4 did not exhibit strong accumulation of H3K27me3 even in WT, suggesting that H3K27 methyltransferases may not directly regulate CTCF binding in the Class 4 (Fig. 1F). Additionally, in “Increased” Class 2 and Class 5, H3K9 methyltransferase deficiency led to an increase in H3K27me3. Despite this increase, CTCF binding in the Class 2 was elevated in TKO, whereas in the Class 5, H3K27me3 increased in 5KO, and CTCF binding increased only after the removal of H3K27me3 (Fig. 1F-H). These findings suggest the existence of additional factors beyond H3K27me3 that regulate the differential behaviour of CTCF binding in the Class 2 and the Class 5. Regarding the “Decreased” class, an increase in H3K9me3 was observed in Class 1 upon *Setdb1* KO, and an increase in H3K9me2 was observed in Class 2 upon TKO (Supplementary Fig. S1D, E). This suggests that the reorganization of H3K9 methylation states due to the loss of each H3K9 methyltransferase inhibits CTCF binding.

Since certain fractions of CTCF binding are sensitive to DNA methylation (Kim et al. 2015), we examined the relationship between “Increased” CTCF binding sites and DNA methylation by reanalyzing the previously reported global DNA methylation status of WT and 5KO iMEFs (Fukuda et al. 2023)(Supplementary Fig. S1F). The results show that significant reduction of DNA methylation is induced in 5KO at “Increased” sites other than class 4. It is reported that the contribution of DNA demethylation to the newly emerged CTCF binding in *Setdb1* KO mESCs is mostly dispensable (Tam et al. 2024). However, the certain fractions of “Increased” sites observed in the H3K9 methyltransferases KO iMEFs may be dependent on DNA demethylation.

### H3KG/K27 methyltransferases inhibit CTCF binding at repetitive sequences

All H3K9 methyltransferases are known to repress transposons for H3K9 methylation (Matsui et al. 2010; Maksakova et al. 2013; Bulut-Karslioglu et al. 2014), and it was reported that SETDB1 and EHMT2 deposits H3K9 methylation on transposons, such as SINE B3, to prevent CTCF binding (Jiang et al. 2020; Gualdrini et al. 2022; Sun et al. 2024; Tam et al. 2024). In addition, H3K27me3 is known to accumulate on transposons in conditions where H3K9 methylation is low, such as in female primordial germ cells (Huang et al. 2021) or H3K9 methyltransferase-deficient cells (Fukuda et al. 2023). These findings implicate that SUV39H1/2 and H3K27 methyltransferases also inhibit CTCF binding at transposons. To explore this, we examined the types of transposons enriched in each class. Consistent with previous reports that SETDB1 inhibits CTCF binding at SINE B3 (Gualdrini et al. 2022; Sun et al. 2024; Tam et al. 2024), we observed a strong accumulation of SINE B3 in “Increased” Class 1 (Fig. 2A). Furthermore, SINE B3 was also enriched in “Increased” Class 2, which is regulated by *Suv3Sh1/2*, and in “Increased” Class 3, which is controlled by *Ehmt1/2*. This suggests that CTCF binding at SINE B3 is suppressed by all H3K9 methyltransferases (Fig. 2A). Although CTCF binding at SINE B3 is inhibited by all H3K9 methyltransferases, most SINE B3 copies lack CTCF binding (Fig. 2B). An analysis of the epigenomic profile of SINE B3 revealed that, compared to the consensus peak, H3K9me3 levels were higher in WT for “Increased” Class 1, 2, and 3 (Fig. 2C). Upon *Setdb1* KO, H3K9me3 levels at SINE B3 decreased in “Increased” Class 1, whereas “Increased” Class 2 remained within regions of high H3K9me3 (Fig. 2C). H3K9me2 was depleted at SINE B3 in WT but was elevated exclusively in “Increased” Class 3 under 5KO conditions (Fig. 2C). These findings indicate that the regulation of H3K9 methylation on SINE B3 is mediated by different methyltransferases depending on the copy of SINE B3. Phylogenetic analysis of SINE B3 showed that each class was not assigned to distinct clusters, suggesting that the regulatory differences among SINE B3 copies are not due to differences in SINE B3 subfamilies (Fig. 2D). Given that compartment patterns differ between classes (Supplementary Fig. S1C), we examined the compartment distribution of SINE B3 and found significant differences: SINE B3 in “Increased” Class 1 was enriched in the A compartment, Class 2 in the B compartment, and Class 3 in an intermediate level (Fig. 2E, F). These findings suggest that SINE B3 undergoes differential CTCF suppression by distinct H3K9 methyltransferases depending on the chromatin compartment in which it resides.

Unlike “Increased” Class 1/2/3, any transposons were enriched in “Increased” Class 4, which is regulated by H3K27 methyltransferases (Fig. 2A). However, various transposons including L1 and ERV were enriched in “Increased” Class 5, which is controlled by both H3K9 and H3K27 methyltransferases (Fig. 2A). These findings suggest that H3K9 and H3K27 methylation redundantly suppress potential CTCF binding sites within various transposons. We observed that H3K27me3 was redistributed to transposons enriched in “Increased” Class 5 (IAPLTR2_Mm, L1Fd_2) under 5KO conditions (Fig. 2G, Supplementary Fig. S2A). This suggests that the redistribution of H3K27me3 to transposons following the loss of H3K9 methylation plays a role in preventing CTCF binding at these transposon sites.

**Figure 2.**
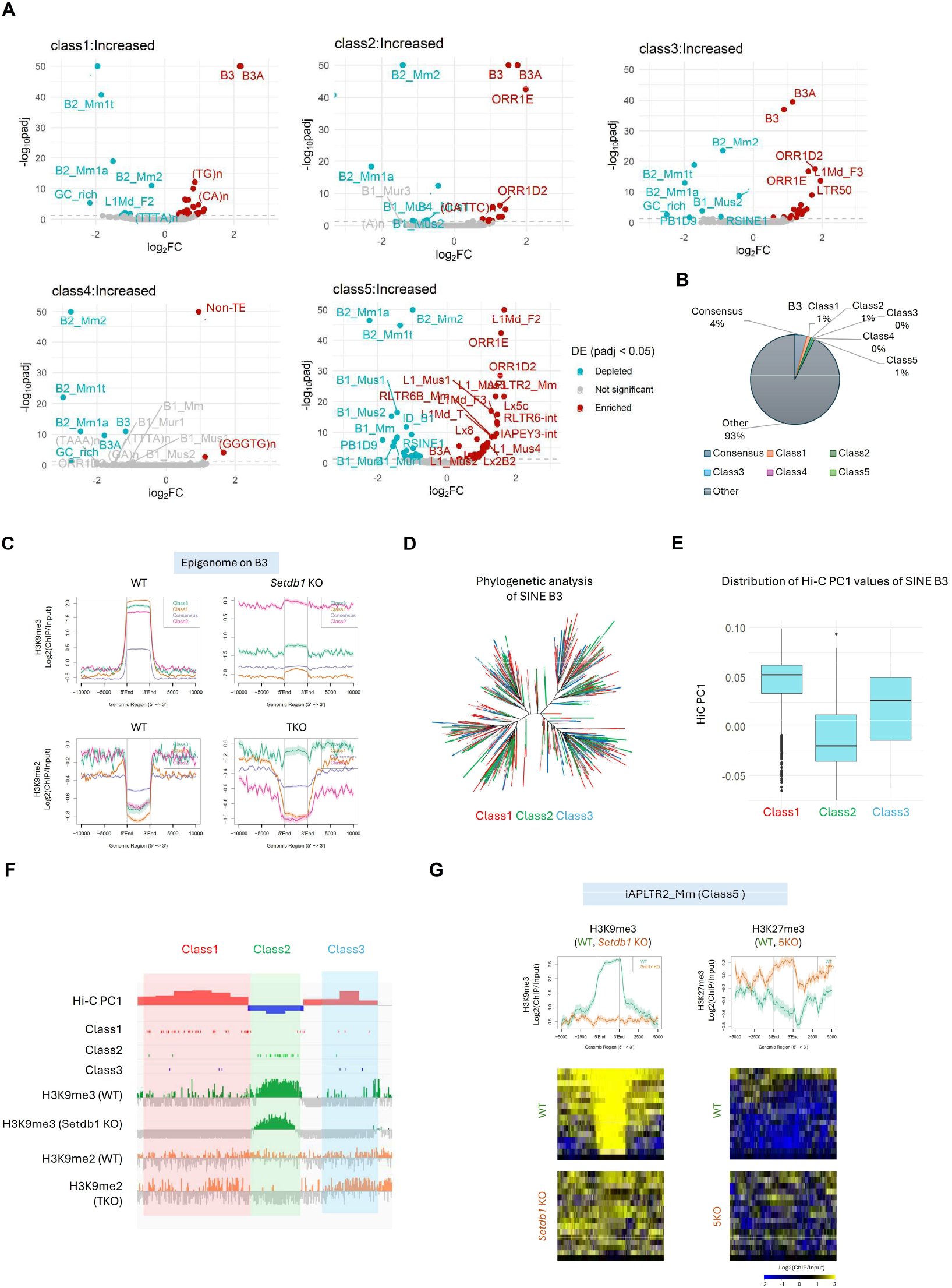
Prevention of abnormal CTCF bindings at repetitive elements by H3K9/K27 methylation. (A) Enrichment of repeat type in increased DE peaks. The volcano plots represent the enrichment of each repeat type in DE peaks, with the x-axis showing Log_2_(FC) and the y-axis showing -log_10_(adjusted P-value). Repeat types that are significantly enriched or depleted (adj. P-value < 0.01) are highlighted in red and blue, respectively. (B) Fraction of SINE B3 transposons overlapping with “Increased” DE peaks. (C) Enrichment of H3K9 methylation around SINE B3 transposons overlapping with “Increased” DE peaks. (D) Phylogenetic analysis of SINE B3 transposons. SINE B3 transposons overlapping with “Increased” DE peaks are color-coded according to their respective classes. (E) The distribution of Hi-C PC1 values in genomic regions where SINE B3 transposons overlap with “Increased” DE peaks. (F) Representative genomic regions where SINE B3 transposons overlap with “Increased” DE peaks. SINE B3 elements overlapping with Class 1 are predominantly found in strong A compartments, those overlapping with Class 2 are mainly in B compartments, and those in Class 3 are frequently located in intermediate regions between Class1 and Class2. (G) Epigenome profiles around IAPLTR2_Mm elements overlapping with “Increased” Class5. In WT iMEFs, H3K9me3, regulated by SETDB1, is enriched. However, when H3K9 methylation is lost, H3K27me3 becomes enriched instead.

Unlike the “Increased” Class, the “Decreased” Class showed only a weak accumulation of a limited number of transposons, such as the IAPEY family in Class 1 (Supplementary Fig. S2B). Interestingly, GC-rich elements and simple repeats with high GC content were strongly enriched, particularly in “Decreased” Class 3, 4, and 5 (Supplementary Fig. S2B). Since GC-rich regions are often found in promoter regions, this enrichment aligns with the fact that these classes are frequently located in promoters (Supplementary Fig. S1B). These results indicate that H3K9 and H3K27 methyltransferases function both to promote and inhibit CTCF binding on repetitive elements.

### Association between changes in CTCF binding profiles and alterations in gene expression and insulation

CTCF functions as an insulator, regulating the three-dimensional genome structure and gene expression (Rao et al. 2014). To investigate how the changes in CTCF binding we identified affect gene expression and genome architecture, we reanalyzed our previously conducted RNA-seq and Hi-C data (Fukuda et al. 2023). To analyze the association between CTCF DE peaks and gene expression changes, we identified differentially expressed genes (DE genes) in each sample and calculated the enrichment of DE genes around DE peaks (Fig. 3A, Supplementary Fig. S3A). Specifically, genes upregulated in 5KO and 5KO+DS were highly enriched around “Increased” Class 3 and Class 5, respectively. In contrast, genes downregulated in *Setdb1* KO, TKO, 5KO and 5KO+DS were strongly enriched around “Decreased” Class 1, Class 2, Class 3 and Class 5, respectively (Fig. 3B, Supplementary Fig. S3B). Therefore, changes in CTCF binding caused by the loss or inhibition of H3K9/K27 methyltransferases may alter the expression of nearby genes.

**Figure 3.**
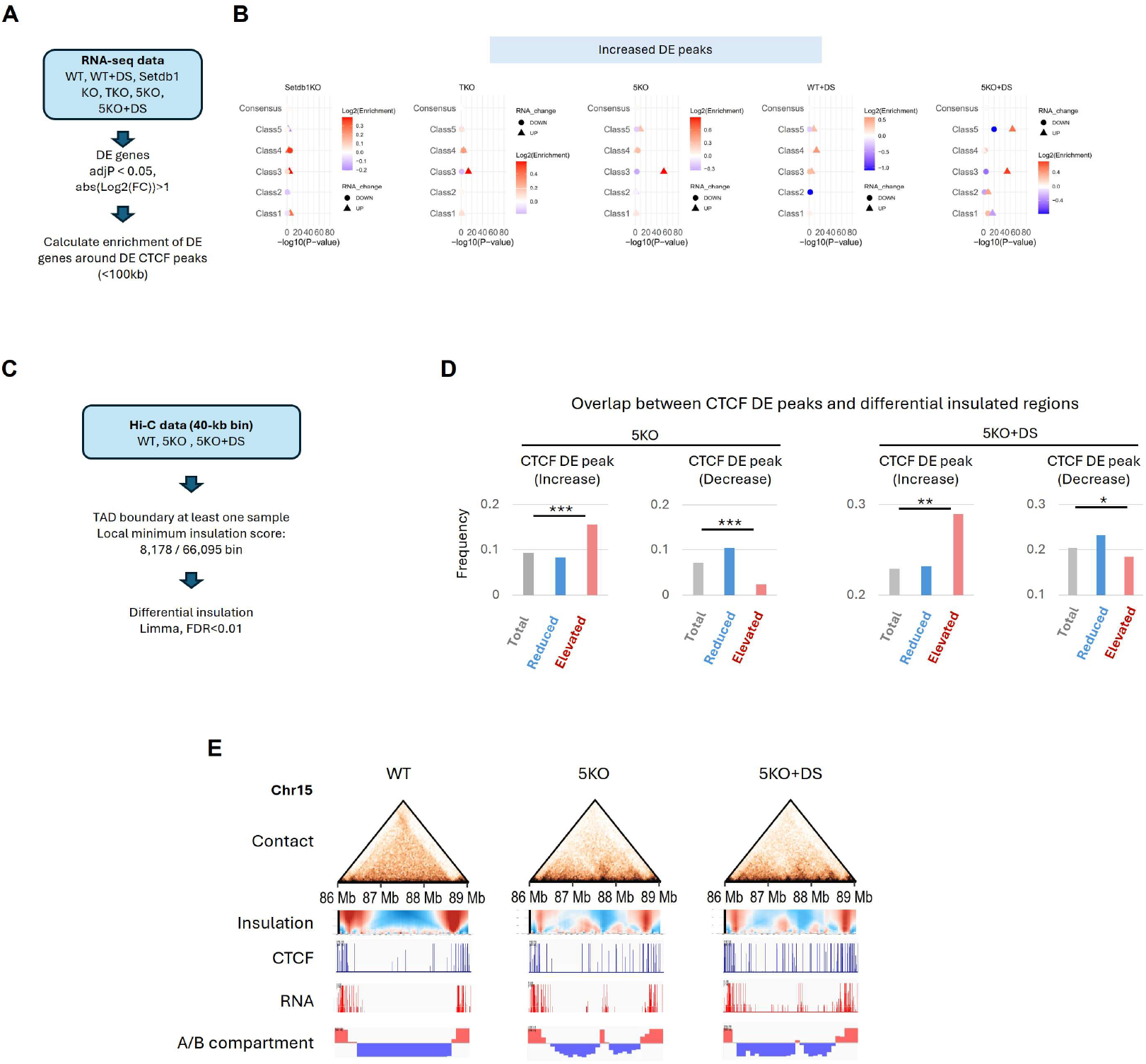
Association of DE peaks with changes in gene expression and 3D genome structure. (A) A process for analyzing the association between DE peaks and changes in gene expression. (B) A dot plot showing the enrichment of differentially expressed genes around DE peaks. The x-axis represents -log_10_(adj. P-value). The enrichment results for upregulated and downregulated genes are represented by circles and triangles, respectively. The degree of enrichment (log_2_(fold change)) is color-coded, with red indicating enrichment and blue indicating depletion triangles and circles, respectively. The degree of enrichment (log_2_(Enrichment)) is color-coded, with red indicating enrichment and blue indicating depletion. (C) A process for analyzing the association between DE peaks and changes in insulation. (D) Frequency of overlap between CTCF DE peaks and differential insulated regions. (E) Representative regions of differential insulation overlapping with “Increased” DE peaks. The figure displays, from top to bottom, the Contact heatmap, insulation map, CTCF ChIP-seq signal, RNA-seq signal, and compartment information. In this region, an increase in CTCF binding, the emergence of new insulation sites, an increase in gene expression, and a B-to-A compartment change are observed in both 5KO and 5KO+DS iMEFs.

The association of both upregulated and downregulated genes with CTCF binding changes in 5KO and 5KO+DS prompted us to investigate insulation changes in these cells. Using our previously reported Hi-C data from WT, 5KO, and 5KO+DS (Fukuda et al. 2023), we extracted insulated regions at 40-kb resolution and identified regions where insulation changed in 5KO and 5KO+DS (Fig. 3C). We detected 472 and 523 regions with increased insulation and 1,477 and 1,073 regions with decreased insulation in 5KO and 5KO+DS, respectively (Supplementary Fig. 3C). In both 5KO and 5KO+DS, regions with elevated insulation significantly overlapped with “Increased” CTCF peaks, while regions with reduced insulation significantly overlapped with “Decreased” CTCF peaks (Fig. 3D). Furthermore, some of these CTCF binding and insulation changes were also associated with gene expression changes (Fig. 3E). These results suggest that H3K9 methylation and H3K27 methylation function independently or cooperatively to maintain the homeostasis of CTCF binding, 3D genome structure, and gene expression.

### Prevention of CTCF by H3KG methylation in early embryo specific CTCF binding sites

The epigenome and 3D genome structure undergo dynamic changes during development. Recently, an analysis of CTCF binding dynamics in mouse early embryos reported that cleavage stage-specific CTCF binding sites are derived from the SINE B2 and B3 families (Wang et al. 2024).

These cleavage stage-specific CTCF binding sites acquire H3K9me3 during development, leading to the loss of CTCF binding (Wang et al. 2024). This report prompted us to compare CTCF DE peaks, which we identified, with the CTCF binding and epigenome profiles during early development using publicly available data (Liu et al. 2016; Wang et al. 2018; Wang et al. 2024). The CTCF binding levels at “Increased” DE peaks dynamically fluctuate during early development. The CTCF enrichment in “Increased” classes except for Class5 are low in MII oocytes, peak at the 2-cell stage, and then gradually decrease (Supplementary Fig. S4A). In contrast, H3K9me3 follows the opposite pattern, being lowest at the 2-cell stage and increasing thereafter (Supplementary Fig. S4B). The CTCF binding levels of “Increased” Class 5 remained low throughout development (Supplementary Fig. S4A). Since the global levels of CTCF and H3K9me3 fluctuate significantly during development, we set CTCF Stable Peaks as internal control regions, where CTCF binding remains unchanged across all iMEF conditions analyzed. The CTCF/H3K9me3 levels in each developmental stage were then normalized using the median value of the Stable peaks. Additionally, since CTCF-mediated chromatin loops emerge from the 8-cell stage onward (Wang et al. 2024), we focused our analysis on the 8-cell stage and Inner Cell Mass (ICM). When comparing CTCF binding in “Increased” CTCF peaks between the 8-cell stage and ICM, particularly in Class 1, CTCF binding at the 8-cell stage was significantly higher than that in ICM (Fig. 4A). On the other hand, H3K9me3 was notably lower at the 8-cell stage than in ICM, especially in Class 1 (Fig. 4B). A similar trend was observed in Class 2 and Class 3, though the degree was not as pronounced as in Class 1 (Fig. 4A, B). CTCF/H3K9me3 in “Increased” Class 4 and Class 5 showed no or little change between the 8-cell stage and ICM (Fig. 4A, B). These results suggest that CTCF binding sites inhibited only by H3K9 methylation become low in H3K9me3 at the cleavage stage, allowing CTCF binding in its place. Notably, CTCF binding sites regulated by *Setdb1* are particularly associated with cleavage-stage-specific CTCF binding profiles. To investigate whether *Setdb1*-dependent inhibited CTCF binding sites are involved in the formation of early embryo-specific 3D genome structures, we compared the insulation status between the 8-cell stage and ICM. The results showed that “Increased” Class 1 specifically functions as an insulator at the 8-cell stage (Fig. 4C, D). In consistent with low CTCF binding throughout early development in “Increased” Class 5, Class 5 does not show insulation function at the 8-cell stage (Supplementary Fig. S4C). When examining the H3K27me3 profile at the 8-cell stage, Class 5 exhibited particularly high levels of H3K27me3, embedding within early embryo-specific H3K27me3 domains (Fig. 4E, F). These findings suggest that H3K27me3 may prevent abnormal CTCF binding during early embryo development, when H3K9 methylation becomes unstable.

**Figure 4.**
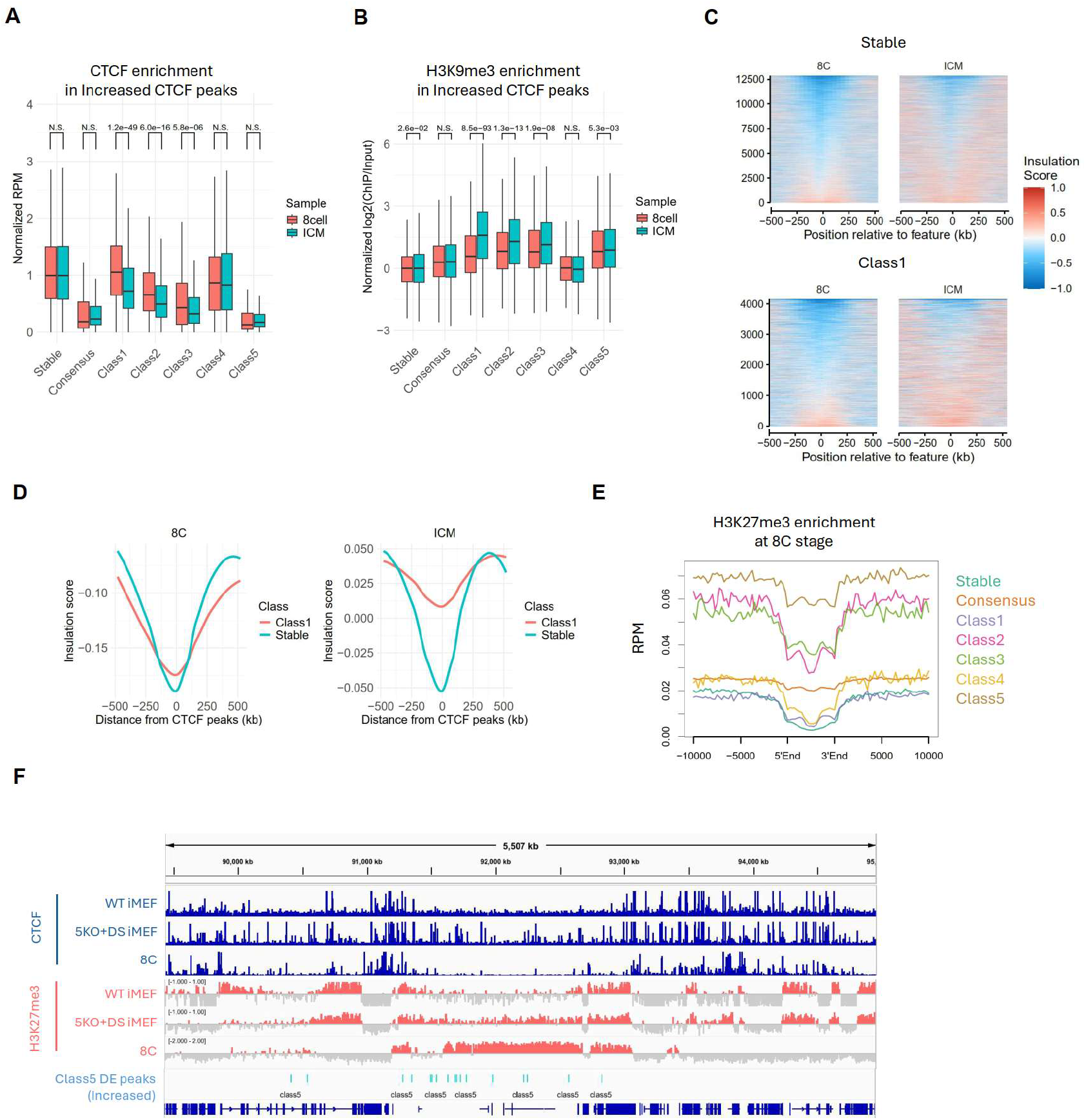
Association of DE peaks with 3D genome structures in mouse early embryos. (A) CTCF accumulation in “Increased” DE peaks in early mouse embryos. The Y-axis represents CTCF accumulation, corrected by the median value of Stable CTCF peaks in each developmental stage. CTCF accumulation in DE peaks is tested between the 8-cell stage and ICM using a t-test. (B) H3K9me3 accumulation in “Increased” DE peaks in early mouse embryos. The Y-axis represents H3K9me3 accumulation (log_2_(ChIP/Input)), corrected by the median value of Stable CTCF peaks in each developmental stage. H3K9me3 accumulation in DE peaks is tested between the 8-cell stage and ICM using a t-test. (C, D) A Tornado plot showing the insulation status in Stable and “Increased” Class1 DE peaks in 8 cells and ICM. Fig. 4C displays the results for each region in a heatmap, while Fig. 4D shows the average values of regions. The X-axis represents the distance from CTCF peaks. (E) Enrichment of H3K27me3 in 8-cell stage around “Increased” DE peaks. Class 5 is embedded in regions with high H3K27me3 enrichment. (F) Representative regions of “Increased” Class5 DE peaks embedded in H3K27me3 enriched regions in 8-cell stage.

## Discussion

In this study, we utilized various H3K9 methyltransferase-deficient cells and H3K27 methyltransferase EZh1/2 inhibitor to elucidate the specificity and redundancy of each enzyme in the regulation of CTCF. We demonstrated that both types of methylation play a role in preventing abnormal 3D genome architecture and transcription by inhibiting CTCF binding. Loss of both H3K9 and H3K27 methylation resulted in dramatic changes in the CTCF binding profile, indicating that these modifications function redundantly to maintain the robustness of the CTCF binding profile and 3D genome organization. Furthermore, classification of CTCF binding changes under different cellular conditions enabled us to identify the characteristics of genomic regions where each enzyme exhibits specificity or functions cooperatively. CTCF primarily peaks in the A compartment and is excluded from the B compartment. Consequently, the B compartment forms a large single TAD, where sub-TAD structures are difficult to establish. However, the underlying mechanism was unclear. Our analysis of cells deficient in both H3K9 and H3K27 methylation demonstrated that these modifications inhibit CTCF binding within the B compartment and prevent the formation of sub-TAD structures in the B compartments.

Transposons, which occupy about half of the mammalian genomes, can serve as major sources of binding sites for various transcription factors, but they are often covered by repressive chromatin modifications. It has been previously known that H3K9 methylation regulates CTCF binding at transposons. However, our analysis revealed the presence of additional potential CTCF binding sites that are repressed by both H3K27me3 and H3K9 methylation. Furthermore, we found that not only the previously focused SINE B2/B3 elements but also a wide variety of transposons can potentially function as CTCF binding sites. Recently, we reported that in the absence of both H3K9 methylation and H3K27me3, uH2A functions to maintain compartmentalization (Fukuda et al. 2025). This raises the possibility that uH2A may also mask additional potential CTCF binding sites.

The state of H3K9 methylation dynamically changes during early development, and accordingly, genomic regions that usually do not bind CTCF, including transposons, can become specifically bound by CTCF in early embryos (Wang et al. 2024). We revealed that CTCF binding sites that are repressed by H3K9 methyltransferases in iMEFs show decreased H3K9 methylation and increased CTCF binding in early embryos. Thus, H3K9 methyltransferases prevent the formation of an early embryo-like 3D genome architecture in somatic cells. On the other hand, CTCF binding sites that are inhibited by both H3K9 methylation and H3K27me3 in iMEFs form H3K27me3 domains in early embryos, resulting in consistently low CTCF binding throughout early development. Such regions are predominantly found in the B compartment, where abnormalities in the 3D genome are robustly suppressed. Such domain formation of H3K27me3 is also observed in aging cells, suggesting that H3K27 methylation contributes to the maintenance of 3D genome architecture even when H3K9 methylation is reduced.

The molecular mechanism by which H3K27me3 inhibits CTCF binding remains unclear. While DNA methylation is known to suppress CTCF binding, DNA methylation and H3K27me3 are anti-correlated, and DNA methylation levels decrease in regions where H3K27me3 infiltrates in 5KO cells (Fukuda et al. 2023). “Increased” Class5 DE peaks already showed strong reduction of DNA methylation in 5KO iMEFs compared to WT iMEFs (Supplementary Fig. S1F). Therefore, it is unlikely that DNA methylation is responsible for inhibiting CTCF binding in H3K27me3-rich regions. Further studies are needed to elucidate the mechanism by which H3K27me3 suppresses CTCF binding.

In conclusion, this study revealed previously unidentified potential CTCF binding sites and proposed a new regulatory layer that maintains the CTCF binding profile. Our findings provide new insights into the mechanisms controlling 3D genome architecture and gene expression regulation, shedding light on how these mechanisms contribute to cellular development and tissue-specific regulation.

## MATERIALS AND METHODS

### Cell culture

iMEFs were maintained in Dulbecco’s modified Eagle’s medium (Nacalai tesque, 08458-16) containing 10% fetal bovine serum (Biosera, FB1061), MEM Non-Essential medium and 2-Mercaptoethanol (Nacalai tesque, 21417-52). To inhibit EZH1/2 catalytic activity, iMEFs were cultured for seven days with 1 µM DS3201 (DS).

### CTCF ChIP-seq

5 × 10^5^ iMEFs were crosslinked in 1 mL PBS supplemented with 0.1 mL Crosslinking Buffer (100 mM NaCl, 1 mM EDTA, 0.5 mM EGTA, 50 mM HEPES [pH 8.0], 11% PFA) at 25°C for 20 min. The reaction was quenched by adding 0.125 M glycine. After washing with PBS, the pellet was resuspended in 300 µL SDS lysis buffer (50 mM Tris-HCl [pH 8.0], 10 mM EDTA, 1% SDS, protease inhibitor). Chromatin was sonicated using a Picoruptor (Diagenode) with a cycle of 30 sec on/30 sec off for a total of five cycles. The lysate was then centrifuged at 15,000 rpm for 10 min at 4°C. The supernatant was diluted with 1.2 mL Chromatin Dilution Buffer (10% Tris-HCl [pH 7.5], 140 mM NaCl, 1 mM EDTA, 0.5 mM EGTA, 1% Triton X-100, 0.1% sodium deoxycholate, 1× cOmplete EDTA-free protease inhibitor (Roche, 11836170001), 1 mM PMSF). From the diluted chromatin solution, 100 µL was set aside for input control, and the remaining portion was incubated with 8 µg of anti-CTCF antibody (Active Motif #61311) and 10 µl of Protein A/G agarose beads (Santa Cruz, sc-2003). The mixture was rotated at 4°C overnight. After rotating the solution, the beads were sequentially washed by low salt buffer (150 mM NaCl, Tris-HCl pH8.0, 2 mM EDTA, 1% Triton X-100, 0.1% SDS), high salt buffer (500 mM NaCl, Tris-HCl pH8.0, 2 mM EDTA, 1% Triton X-100, 0.1% SDS), LiCl buffer (25 mM LiCl, 10 mM Tris-HCl pH8.0, 1 mM EDTA, 1% NP40, 1% DOC), TE, and were suspended in Salting method buffer (20 mM Tris pH8.0, 10mM EDTA, 400 mM NaCl). Input and ChIP samples were de-crosslinked by RNaseA, SDS and Protease K treatment, and purified by QIAquick PCR purification kit (Qiagen).

### Preparation of ChIP-seq library

The ChIP DNA was fragmented by Picoruptor (Diagenode) for 10 cycles of 30 seconds on, 30 seconds off. Then, ChIP library was constructed by KAPA Hyper Prep Kit (KAPA BIOSYSTEMS) and SeqCap Adapter Kit A (Roche) according to manufacturer instructions. The concentration of the ChIP-seq library was quantified by KAPA Library quantification kit (KAPA BIOSYSTEMS). ChIP sequencing was performed on a HiSeq X platform (Illumina). We performed two biological replicates for ChIP-seq.

### Hi-C data analysis

Hi-C data processing was done by using Docker for 4DN Hi-C pipeline (v43, https://github.com/4dn-dcic/docker-4dn-hic). The pipeline includes alignment (using the mouse genome, mm10) and filtering steps. After filtering valid Hi-C alignments, .*hic* format Hi-C matrix files were generated by Juicer Tools (Durand et al. 2016) using the reads with MAPQ>10. The A/B compartment (compartment score) profiles (in 250 kb bins) in each chromosome (without sex chromosome) were calculated from .*hic* format Hi-C matrix files (intrachromosomal KR normalized Hi-C maps) by Juicer Tools (Durand et al. 2016) as previously described (Miura et al. 2018). We averaged Hi-C PC1 values in each 250 kb bin from two biological replicates for the downstream analysis. We used GENOVA tools (van der Weide et al. 2021) for producing aggregate TAD plots, insulation analysis and pyramid plots.

The insulation score was calculated for each 40-kb bin using GENOVA tools. For each 40-kb bin, the local minimum of the insulation score was identified within a ±200-kb window (upstream 200 kb and downstream 200 kb). Bins with local minima were assigned as TAD boundaries. Out of 66,095 bins, 8,178 were identified as TAD boundaries in at least one sample. Next, the insulation scores of all identified TAD boundaries were compared between WT and other conditions. Regions with significant changes in insulation (differential insulation) were identified using Limma (FDR < 0.05).

### ChIP-seq analysis

CTCF ChIP-seq analysis was performed using Nextflow with the nf-core/chipseq pipeline (version 2.0.0) (Ewels et al. 2020). The analysis was conducted using the mouse mm10 genome as the reference. Differential peaks were identified using *diffBind* (FDR<0.05) (Ross-Innes et al. 2012) based on narrow peaks called by MACS2 (Zhang et al. 2008) in the Nextflow pipeline. The commonality of differential peaks across samples was analyzed using bedtools (Quinlan and Hall 2010), and peaks were classified into five categories accordingly.

### RNA-seq analysis

Adaptor sequences and low-quality bases in reads were trimmed using Trim Galore version 0.3.7 (http://www.bioinformatics.babraham.ac.uk/projects/trim_galore/). To calculate RPM values in each 250 kb bin, trimmed reads were aligned to the mm10 genome build using bowtie version 0.12.7 with –m 1. Duplicated reads were removed using samtools version 0.1.18. The number of mapped reads in each 250 kb bin was counted by featureCounts (Liao et al. 2014), then, RPM values in each 250 kb bin was calculated. To identify differentially expressed genes or repeats, the trimmed reads were mapped to the mouse GRCm38 genome assembly using TopHat (v2.1.1) with –g 1 (Trapnell et al. 2009). After read mapping, the number of reads mapped in genes or repeats was counted by TEtranscripts (v1.4.11) with default parameters) (Jin et al. 2015). We performed two biological replicates for RNA-seq and identified DE genes by DESeq2 (adj. P-value < 0.05, FC > 2) (Love et al. 2014).

### Visualization of NGS data

The Integrative Genomics Viewer (IGV) (Robinson et al. 2011) was used to visualize NGS data. For Hi-C contact matrix and correlation matrix, we used Juicer Tools (Durand et al. 2016).

### Statistical analysis

All methods for statistical analysis and P-values in this study are described in each figure legend and included in each figure, respectively.

### Data availability

All NGS data used in this study are described in Supplementary Table S1.

## ACKNOWLEDGEMENTS

We thank the staff of the Support Unit for Bio-Material Analysis (BMA) at the RIKEN Center for Brain Science (CBS) Research Resources Division (RRD) for NGS library construction, DNA sequencing and flow cytometry. We would also like to thank our colleagues at Shinkai laboratory for their support and valuable comments.

## Author contributions

K.F. and Y.S. designed and conceived the study. K.F. and Y.S. supervised the study and interpreted the data. C.S. performed molecular and cellular experiments and generated the ChIP-seq, RNA-seq and Hi-C-seq libraries. K.F. performed informatics analysis of generated NGS data. K.F. and Y.S. wrote the manuscript and prepared figures. All authors read, discussed, and approved the manuscript.

## FUNDING

RIKEN internal research fund (Pioneering project ‘Genome building from TADs’) (to Y.S.); Y.S. was also supported by the Japan Society for the Promotion of Science (JSPS) [for Grant-in-Aid for Scientific Research [A], JP22H00413; Grant-in-Aid for Scientific Research on Innovative Areas (Research in a proposed research area), JP18H05530]; F.K. was supported by the JSPS [for Grant-in-Aid for Early-Career Scientists, 22K15044]. Funding for open access charge: Japan Society for the Promotion of Science.

## Conflict of interest statement

None declared.

## Notes

### Competing Interest Statement

The authors have declared no competing interest.

### Summary of Updates

The sections Acknowledgements, Funding, and Conflict of Interest have been added to the new version.

