## Supplemental Figure 1 for "H3K27 and H3K9 Methylation Mask Potential CTCF Binding Sites to Maintain 3D Genome Integrity" (from another version of this preprint: version 1, 2025-04-03)

### Supplementary Figure S1

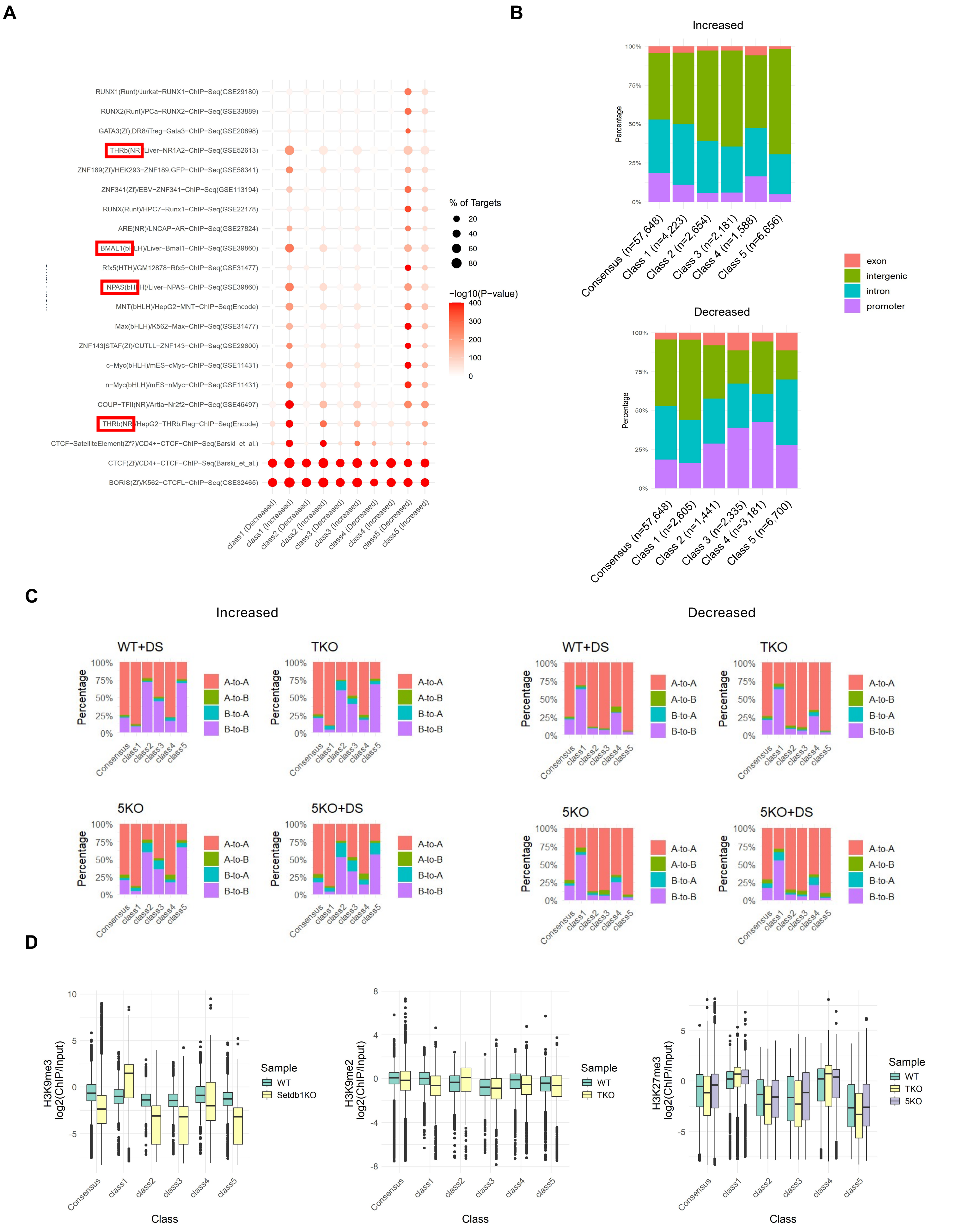

**E**

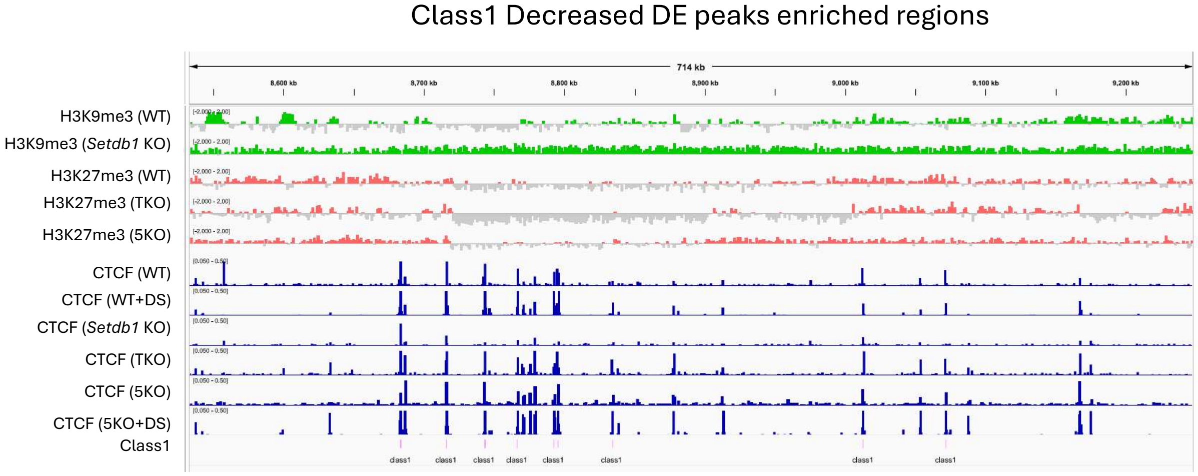


**F**

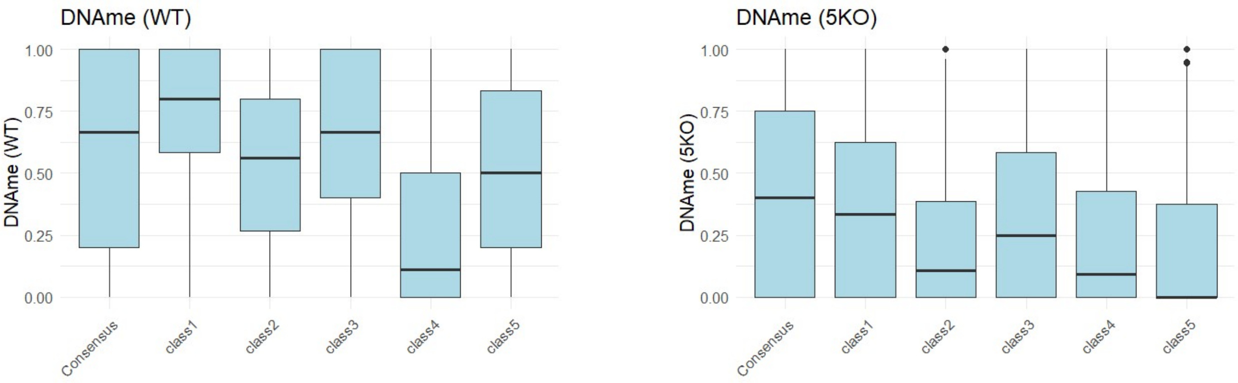


**Supplementary Figure 1. Dysregulation of CTCF binding profiles by H3K9/K27 methylation deficiency.** (A) Motif enrichment in DE peaks. (B) The distribution of DE peaks across genomic regions, including promoters, exons, introns, and intergenic regions. (C) A bar graph showing the frequency of compartment changes in DE peaks. (D) Epigenomic profiles at "Decreased" DE peaks. H3K9me3 enrichment in WT and *Setdb1* KO iMEFs (Top), H3K9me2 enrichment in WT and TKO iMEFs (Middle), and H3K27me3 enrichment in WT, TKO, and 5KO iMEFs (Bottom). (E) Representative region of "Decreased" Class1 enriched regions. (F) Boxplots of DNA methylation levels of "Increased" DE peaks in WT (left) and 5KO (right).
