## Supplemental Figure 4 for "H3K27 and H3K9 Methylation Mask Potential CTCF Binding Sites to Maintain 3D Genome Integrity" (from another version of this preprint: version 1, 2025-04-03)

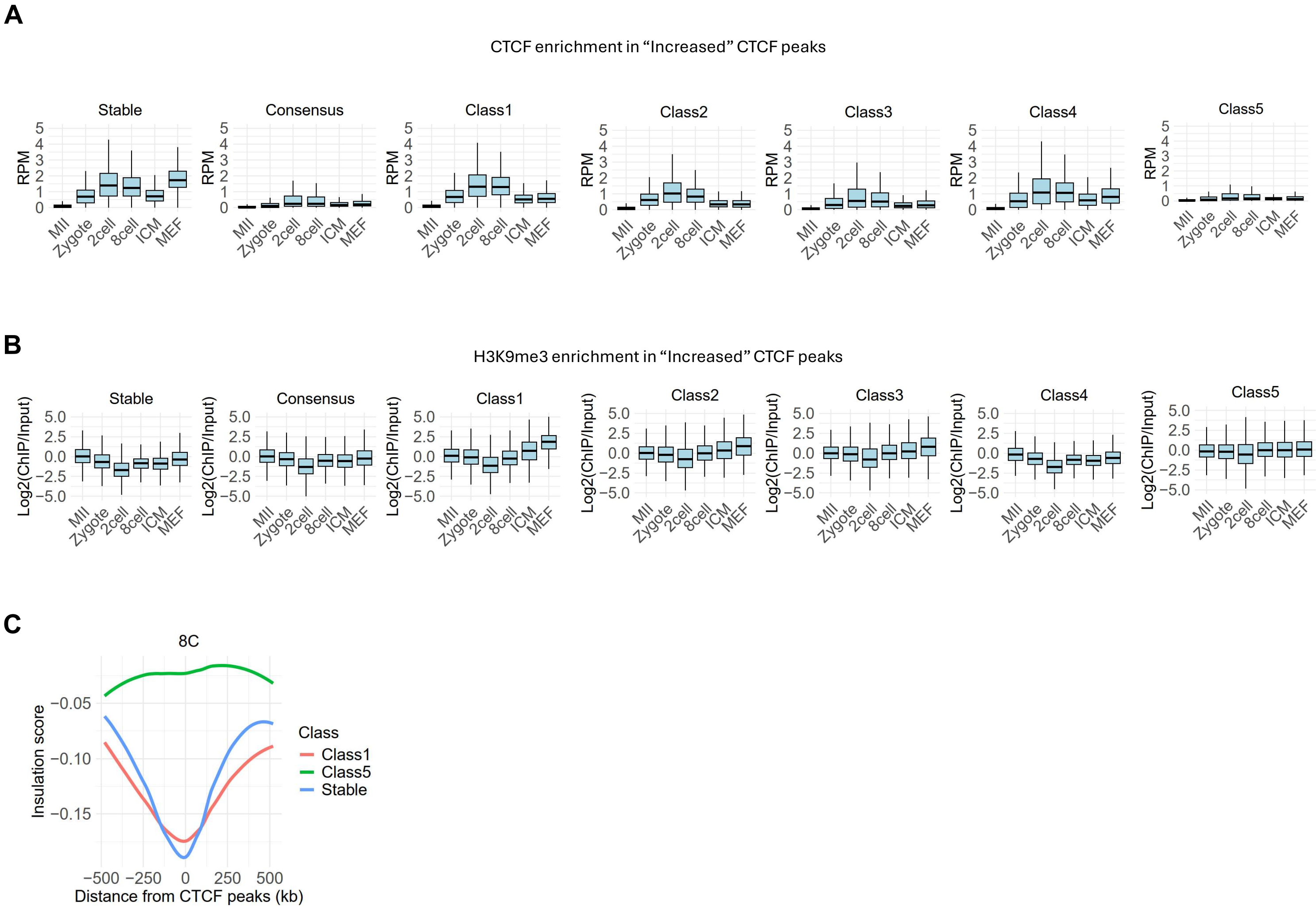


**Supplementary Figure 4. Association of DE peaks with 3D genome structures in mouse early embryos.** (A) Dynamics of CTCF enrichment in Increased DE peaks during early mouse development. (B) Dynamics of H3K9me3 enrichment in Increased DE peaks during early mouse development. (C) Average insulation score in 8-cell stage around Increased DE peaks.
